# Ten-year projection of white-nose syndrome disease dynamics at the southern leading-edge of infection in North America

**DOI:** 10.1101/2020.11.12.379271

**Authors:** Melissa B. Meierhofer, Thomas M. Lilley, Lasse Ruokolainen, Joseph S. Johnson, Steven Parratt, Michael L. Morrison, Brian L. Pierce, Jonah W. Evans, Jani Anttila

**Author notes:** **Corresponding author:** Thomas M. Lilley, Finnish Museum of Natural History, University of Helsinki, Pohjoinen Rautatiekatu 13, 00100 Helsinki, Finland. **Article impact statement:** A projected decline in bat abundance in southern North America.

## Abstract

Predicting the emergence and spread of infectious diseases is critical for effective conservation of biodiversity. White-nose syndrome (WNS), an emerging infectious disease of bats, has resulted in high mortality in eastern North America. Because the fungal causative agent *Pseudogymnoascus destructans* is constrained by temperature and humidity, spread dynamics may vary greatly by geography. Environmental conditions in the southern part of the continent, where disease dynamics are typically studied, making it difficult to predict how the disease will manifest. Herein, we modeled the spread of WNS in Texas based on available cave densities and average dispersal distances of species occupying these sites, and projected these results out to 10 years. We parameterized a predictive model of WNS epidemiology and its effects on hibernatory bat populations with observed environmental data from bat hibernation sites in Texas. Our model suggests that bat populations in northern Texas will be more affected by WNS mortality than southern Texas. As such, we recommend prioritizing the preservation of large overwintering colonies of bats in north Texas through management actions. Our model further illustrates that infectious disease spread and infectious disease severity can become uncoupled over a gradient of environmental variation. Finally, our results highlight the importance of understanding host, pathogen and environmental conditions in various settings to elucidate what may happen across a breadth of environments.

## Introduction

Emerging infectious diseases (EID) of wildlife are increasing in number and threatening several species with extinction (Hayman et al. 2013; Hoberg & Brooks 2015; Tompkins et al. 2015). Emerging infectious diseases are those newly appearing or rapidly increasing in a population (Morse 2004). EIDs occur when pathogenic or putatively pathogenic organisms in the environment have the opportunity to infect new hosts species or populations. Changing environmental conditions can accelerate this process of host-switching by driving changes in host-species’ distributions and by creating new habitable environmental reservoir pathogens (Morse 1991; 1993). Emergence of infectious diseases are therefore associated with a range of causal factors, including ecosystem alterations and movements of pathogens or vectors (Morse 1995). The spread of EIDs is mediated by differences in host ecology, resulting in various patterns of spatial spread (Smith et al. 2002; LaDeau et al. 2008; Meentemeyer et al. 2011). Therefore, understanding and predicting the spatial structure of future EID epidemics requires integration of both the environmental factors and species-specific ecology that can underpin pathogen contact networks (Parratt et al. 2016).

Modeling the spread of EIDs can guide and improve upon effective, science-based management and conservation practices and efforts by identifying key factors driving pathogen persistence and disease dynamics (Keeling & Rohani 2007; Perez & Dragicevic 2009). Although predictive power depends on the accuracy of the data used, models can provide predictive information to target conservation and control measures where the pathogen exists and plan adaptive management strategies where the pathogen may spread (Keeling & Rohani 2007; Cunniffe et al. 2016). These targeted measures can be used to improve the efficacy of conservation measures and help to eradicate infections from the population, protecting host biodiversity (Keeling & Rohani 2007).

An EID of bats known as white-nose syndrome (WNS) threatens the survival of populations of several cave-hibernating species in North America (Gargas et al. 2009; Lorch et al. 2011; Warnecke et al. 2012). Since the first documentation in North America in winter 2006– 2007, the fungal causative agent *Pseudogymnoascus destructans* has spread at a rate of 200 to 900 km per year, and is associated with mortality in excess of 90% (Gargas et al. 2009; Foley et al. 2011; Ingersoll et al. 2013). The first documentation of the disease occurred in Howes Cave near Albany, New York in February 2006, and it has since been documented across a substantial portion of North America (Blehert et al. 2009; Turner & Reeder 2009; Lorch et al. 2016). Although, bat-to-bat transmission is the primary mode of disease dispersal (Raudabaugh & Miller 2013), *P*. *destructans* can persist in an environment devoid of bats (Minnis & Linder 2013; Hoyt et al. 2015; Leopardi et al. 2015). The disease disrupts hibernation behavior through multiple pathways (Verant et al. 2014; Field et al. 2015; Lilley et al. 2017) leading to an increased arousal frequency and ultimately, the depletion of fat reserves (Reeder 2012). This has generated predictions of local extirpations and extinctions of once common bat species (Ingersoll et al. 2016; Pettit & O’Keefe 2017; Frank et al. 2019). There is therefore a need to understand future spread so that conservation efforts can be best prioritized. Moving towards this understanding will require study of how factors associated with WNS transmission work together to influence spread.

The vegetative growth of *P. destructans* is constrained by temperature and humidity (Verant et al. 2012; Marroquin et al. 2017), hence certain environments may not be optimal for growth and thus impede spread. Factors known to be associated with the transmission of the fungus include bat species composition and abundance, population composition (Lilley et al. 2020), geography (e.g., connectivity of hibernacula), and climate (Wilder et al. 2011; Langwig et al. 2012; Maher et al. 2012; Lilley et al. 2018). As a result, the dynamics of WNS disease spread may vary greatly by geography and demography. Predictive modelling of WNS has focused on northeastern United States (e.g., Flory et al. 2012), or used data from that region to parameterize their disease-spread models in other regions (e.g., Maher et al. 2012). Consequently, findings from these studies may not reflect regional differences among bat hibernacula (McNab 1974; Humphries et al. 2002; Brack 2007). It is therefore important to understand and analyze the incidence—and prevalence of—WNS over different spatial and temporal scales to more accurately determine the potential impacts of disease in those regions (Perez & Dragicevic 2009).

Texas provides a unique situation for studying disease spread in that it has the greatest number of bat species of any state in the United States and many of these species were naïve to *P. destructans* until spring 2017, with *P. destructans* documented in the farthest southern locality at 30 parallel north in Texas (Schmidly & Bradley 2016; TPWD 2017, 2019). WNS invasion of Texas is ongoing, with infection first documented during spring 2020 (TPWD 2020). Despite this, minimal information currently exists on whether the environmental conditions in potential bat hibernacula (i.e., caves) (Meierhofer et al. 2019), the spatial layout of caves, and the frequency of suitable caves in the landscape for the persistence of *P. destructans* in the environment in Texas are conducive to the persistence of WNS. It is important to acknowledge that *P. destructans* can exist in the environment without infecting the bat host or causing mortality due to WNS (Lorch et al. 2013b; Hoyt et al. 2015). Unlike the northeast, there has not yet been substantial mortality documented or reported resulting from WNS in Texas. Due to a sense of urgency resulting from the mass declines documented in other regions of North America, researchers are currently deploying treatments in Texas hibernacula as a management strategy to prevent pathogen exposure and to reduce disease severity. Several of these treatments are being deployed in culverts, which are inherently easier to treat in comparison to caves due to their simple design and ease of access. Thus, understanding whether WNS can develop in certain regions in Texas and how the disease will move throughout the southern region is integral in implementing proper conservation and management strategies for caves.

Here, we used demographic and environmental data collected from caves at the leading edge of WNS spread in Texas to parametrize a novel predictive model of fungal epidemiology. To simplify the model and provide robust predictions, our modeling approach does not investigate specific bat species, but combines all species. As such, we refer to a population as all cave-hibernating species combined. Furthermore, using environmental data from caves and PRISM climate data we model the probability of *P. destructans* being able to infect their hosts, leading to symptoms of WNS, and furthermore, the death of the host. Herein, we hypothesized that (1) spread is accelerated by high cave concentrations and bat abundance and (2) disease development is hindered by internal and external environmental conditions affecting both bat ecology and fungal growth. We projected our results 10 years ahead to provide stakeholders information on how the disease will most likely behave to better implement conservation measures.

## Materials and Methods

In Texas where greater than 95% of land is privately owned, caves, as opposed to other hibernacula (e.g., culverts, buildings), are challenging to monitor and manage for WNS because of access restrictions. As such, we focused solely on caves for model development as opposed to other potential hibernacula because of the lack of available data on environmental characteristics of alternative hibernacula across a broad geographic range in Texas and to retain the simplicity of the model.

### Model Development

Our mathematical model is a modification of the model published by Lilley et al. (2018). In comparison with the previously published model, we have simplified the hibernation and transmission dynamics to achieve easier parameterization, and do not consider environmental stochasticity (Supporting Information). The model consists of differential equations, with a periodic temperature forcing, describing the dynamics of the bat hosts and the free pathogenic fungus. We divided the hosts into susceptible, exposed, and infectious, all of which can be either active or hibernating, leading to seven compartments in total. The dynamic state is tracked in a network of patches representing caves within the counties of Texas. We implemented in C++ and full program codes are available at https://github.com/janivaltteri/wnstexas. The full model description is provided in Supporting Information.

To utilize the model on Texas topography, we initially assumed all documented caves could be hibernation sites, and assigned the estimated 4251 hibernation sites obtained from the Texas Speleological Survey database to 94 counties from which we had environmental data. Inside each county (a geographical region used for administrative purposes), we grouped hibernation sites according to cave mean temperatures into bins of 2 °C, following a gaussian distribution with county-specific mean and variance of 3.75 °C. We estimated the mean cave temperature based on a linear model of mean ambient temperature and cave coordinates (Supporting Information). We then used this model to predict mean cave temperatures for the geographic centers of each county. We used the variance of the model residuals to estimate the 3.75 °C variance.

We considered each 2 °C bin of caves as a patch *i* in county *j* with carrying capacity *K*_i,j_ given by the number of hibernation sites in that bin (Fig. 1). We did not assume that all caves were occupied by bats, but rather approximated that 40% of caves were occupied based on our survey data. In total, there were 303 patches. Hibernaculum temperature inside each patch varied sinusoidally with an amplitude estimated for each county (Supporting Information), affecting fungal growth rates inside the hibernaculum. In addition, we assigned each county a mean ambient temperature and annual sinusoidal variation, according to a linear fit (of Fourier coefficients) on the PRISM data.

**Figure 1.**
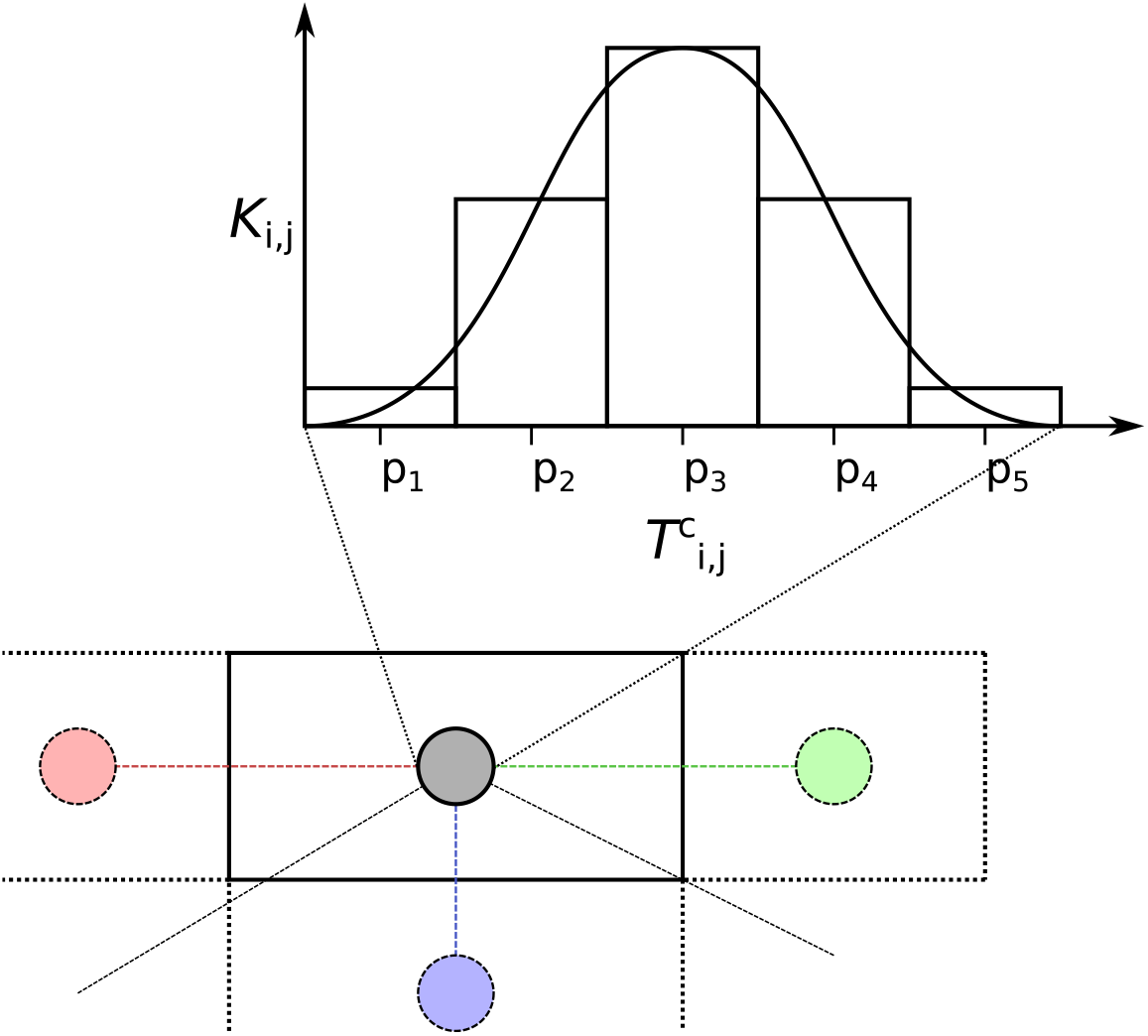
A conceptual drawing of the model spatial setting. The top part shows binning of hibernacula by the within cave mean temperatures according to a Gaussian distribution obtained from a linear model for each county j. Each bin becomes a patch i with a given mean within cave temperature T^{c}_{i,j} and capacity K_{i,j}. In the bottom part, dispersal distances between counties j are the distances between the county midpoints (grey and colored circles).

We implemented patch-to-patch migration (dispersal) as follows: each patch had a fixed proportion of susceptible and exposed bats emigrating per day. To retain simplicity of the model, we do not directly account for species structure variation nor movement among sites during winter. We divided the emigrating bats into recipient patches depending on distance. We assigned a weight 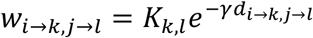, where *d_i→k, j→l_* is the distance from patch *i* in county *j* to patch *k* in county *l* for each connection under a cutoff distance of 100 km, and we calculated the proportion going to a target patch as 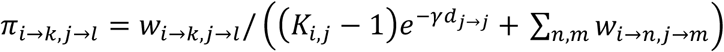. We calculated patch to patch distances *d_j→j_* within a county as mean distances of two randomly placed points inside the county. Distances between counties were simply the distances between the two county midpoints. The parameter γ scales the recipient patch distribution with respect to distance from the focal patch. Under our validated parameter set we used a γ value of 0.00868 km^−1^.

Hibernation strongly affects disease dynamics because *P. destructans* has different effects on active and hibernating bats (Hayman et al. 2016). We determined the duration of hibernation in a patch by ambient and hibernaculum temperatures, *T*_amb_ and *T*_hib_, together with three threshold values α. A patch is in hibernation when either *T_amb_*(*t*) < *α*_*amb*,0_ or *T_amb_*(*t*) < *α*_*amb*,1_; *T_hib_*(*t*) < *α_hib_*. Our parameter set used threshold values α_amb,0_, α_hib_ = 11.5 °C and α_amb,1_ = 12.5 °C (Perry 2013; Meierhofer et al. 2019).

We used a simple linear force of infection in response to environmental fungal density instead of the more sophisticated sigmoidal response used in the model of Lilley et al. (2018). Additionally, we have resorted to simple threshold functions for bat population growth rate and the transfer rates between active and hibernation states. While the original formulation in Lilley et al. (2018) is theoretically sound and results in smoother dynamics, our current formation is analytically more tractable/computationally better suited to integrating real-world variation in parameters. The sigmoidal infectivity response has, however, notable effects on disease dynamics. Therefore, we have replicated all simulation experiments using different sigmoidal parameterizations, and show the resulting effects in Supporting Information.

We visualized model predictions in R with interpolated heat maps generated by the linear bivariate method in the package ‘ akima::interp()’ on an 80×80 grid under default settings. Interpolation predicts values within a convex hull bounding the data points. Therefore, we did not predict beyond the spatial extremes of the data produced by the predictive model described above. We plotted infection as the carrying-capacity scaled predictions from the infection model at 5 and 10 years of simulation, and calculated the loss of bat abundance as the proportional reduction in bats predicted by the infection model relative to a no-infection scenario, obtained by running the model without the fungus and initial infections. We used functions in the ‘Raster’ and ‘ggplot2’ packages to create the figures.

### Model Parameters and Parametrization

We obtained information on number of caves per county within Texas from the Texas Speleological Survey (TSS, https://www.texasspeleologicalsurvey.org/). The TSS, Texas Cave Management Association, local Grottos, biologists, and private landowners provided access to specific cave sites for data collection.

We gathered daily mean ambient temperature data for each county from 1 January 2017 to 31 December 2018 from PRISM (PRISM Climate Group 2018). We used EL-USB-2 Data loggers (Lascar Electronics Inc.) placed within the first third of each cave near roosting bats, when present, to record internal ambient temperature every hour for one year. We deployed loggers at each of 27 caves (13 caves occupied by hibernating bats, 14 unoccupied) distributed in 19 counties across north and central Texas where permission was obtained. We placed loggers near bats or centrally in caves where bats were not present. We obtained information on presence *of P. destructans* within a county from Texas Parks and Wildlife Department (TPWD 2019).

We determined bat and fungal parameter values (Table 1) roughly based on those estimated by Lilley et al. (2018). For our model, we focused solely on using information on bat species known to hibernate in Texas (hibernatory bat populations) as bats that hibernate are affected by WNS. The average number of non-Mexican free-tailed bats (*Tadarida brasiliensis*) was calculated to be 344 based on data collected on bat counts during our previous survey efforts of caves in Texas. We disregarded Mexican free-tailed bat colonies as this species does not tend to hibernate. With our previous survey counts, and documentation of large bat colonies by other researchers (e.g., Tinkle & Milstead 1960; Ammerman et al. 2012; Caire et al. 2019), we approximated that the average number of bats per cave to be 900 for the purpose of our model.

**Table 1.**
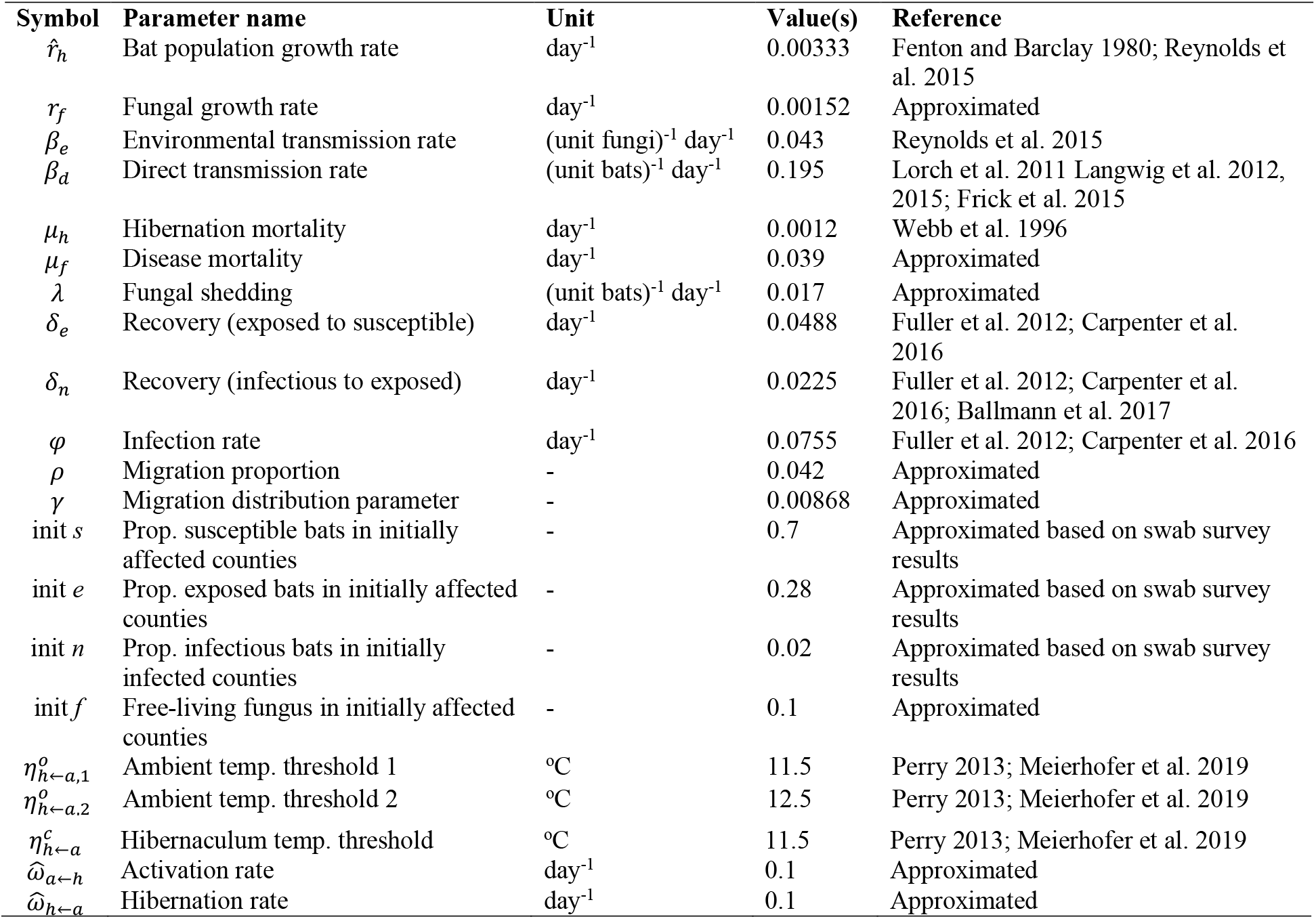
The model parameters and our value estimates after validation step based on 2020 WNS survey data (see Materials and Methods for details). Parameter values are the averaged estimates based on referenced publications and approximated values, and are scaled by the carrying capacity (thus do not directly match values in references).

Direct assignment of parameter values to our model for the actual biological setting would have been challenging because many of the needed values are not known or directly measurable. Instead, we used literature-based values and approximated values of the authors in combination with a parameter estimation step based on the known initial state of the disease in 2018 and survey data from 2020. With the estimation step, we ensured that our parameter set would predict the 2020 observed state from the initial conditions, and thus be in line with the actual known WNS disease dynamics in Texas.

To parameterize the model, we started by constructing a parameter set, which represented our best knowledge of the model parameter values obtained by averaging approximated values of the co-authors (Table 1). We then refined our estimates with an approximate Bayesian computation procedure (Sunnåker et al. 2013). First, we constructed a prior distribution by assigning to each of the parameter values a log-Gaussian distribution with our estimate as the median value and a log-unitary standard deviation, following the reasoning that the true parameter values fall within one order of magnitude from our initial estimate. We then ran 500 simulations with parameters randomly drawn from our prior distributions (Supporting Information) and performed rejection sampling to select appropriate posterior combinations based on WNS 2020 survey data (Fig. 2). With no easy way of assigning likelihood values to our simulations, we used simple rejection thresholds. We used <0.1% disease prevalence in counties where WNS in bats was detected as the rejection criteria, following the reasoning that a small prevalence in bats could already be detected through surveys. We further used <20% free-living fungus prevalence in counties where *P. destructans* was detected as the rejection criteria, because finding fungal growth outside of the bat hosts requires active search of hibernation sites after 2 years of simulation time. Our threshold values were admittedly arbitrary because we had no information on the actual detection effort or efficiency, but these values can easily be improved in future work. Additionally, there were two counties surveyed with neither WNS or *P. destructans* were detected, and we rejected >0.1% disease prevalence and >20% free-living fungus prevalence in these. We then used parameter median values from the accepted combinations (with 5% acceptance rate) as our validated parameter set. We fixed hibernation rate and threshold parameters to our literature-based estimates and were not found by this validation step. We studied the robustness of our results separately in a sensitivity analysis (Supporting Information), where we investigated how varying each parameter by a small increment or decrement changes the simulation outcome in terms of the number of affected patches and reduction in bat numbers.

**Figure 2.**
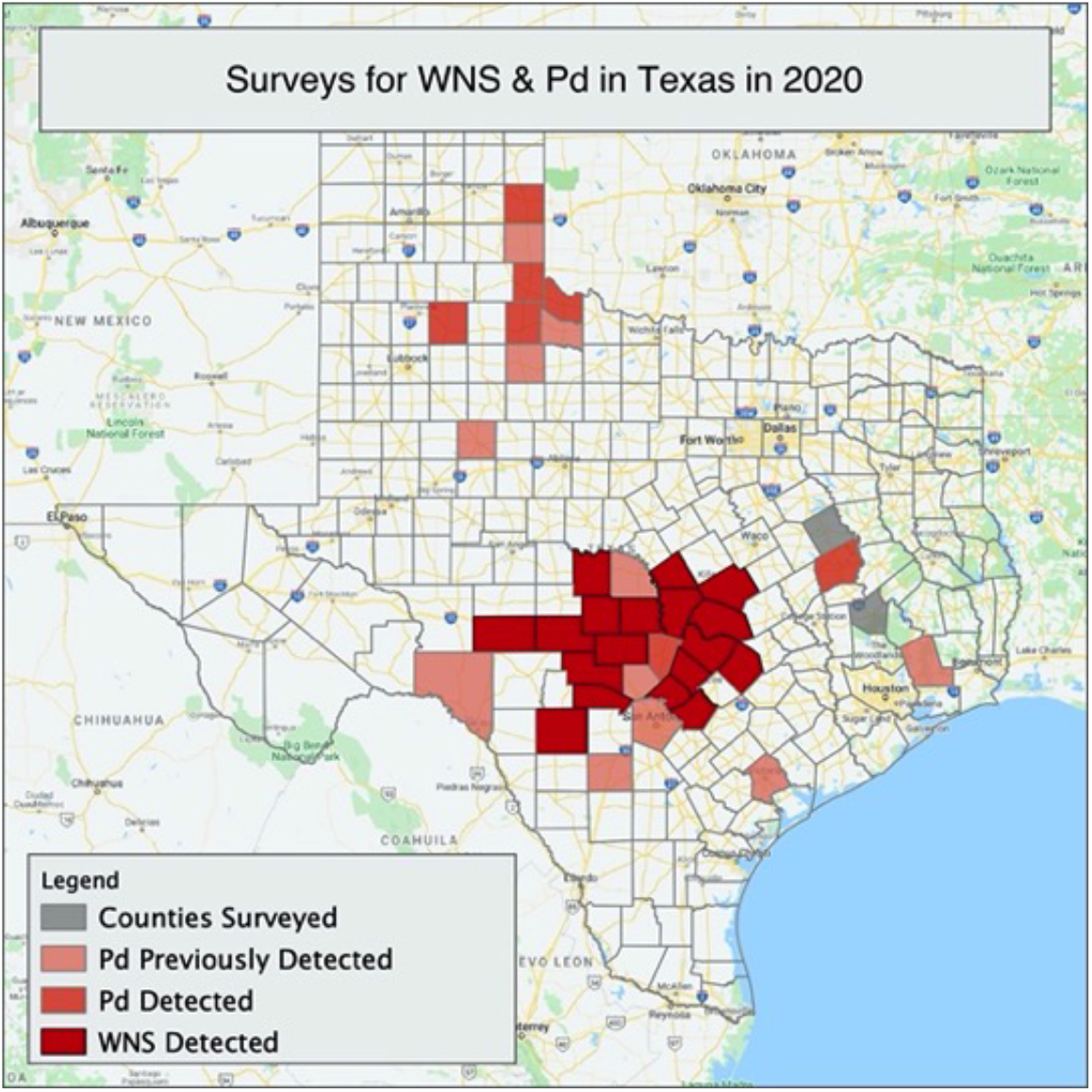
Texas counties where white-nose syndrome (WNS) was detected in 2020 (dark red), where only *P. destructans* was detected in 2020 (medium red), where *P. destructans* was detected in previous years (light red), and counties surveyed in Texas where neither WNS nor *P. destructans* was detected (gray).

## Results

Under our parameter set validated against 2020 WNS survey data, the hibernatory bat population declined 35.6% across 84 counties in 10 years (Fig. 3). After five years, the bat population would be reduced by 19.3% in 70 counties. The simulations did not show local extinctions in any county, but the bat population reduced by 86% (85% after five years) in the most affected site. The most affected counties were in north Texas, with *P. destructans* present at the start of the simulation. The bat population rich mid-Texas are projected to lose between one quarter to half of the bat population (Figs. 3cd, 4a). Density of the free-living fungus and its spores reached high levels in the bat population rich counties (Fig. 4b).

**Figure 3.**
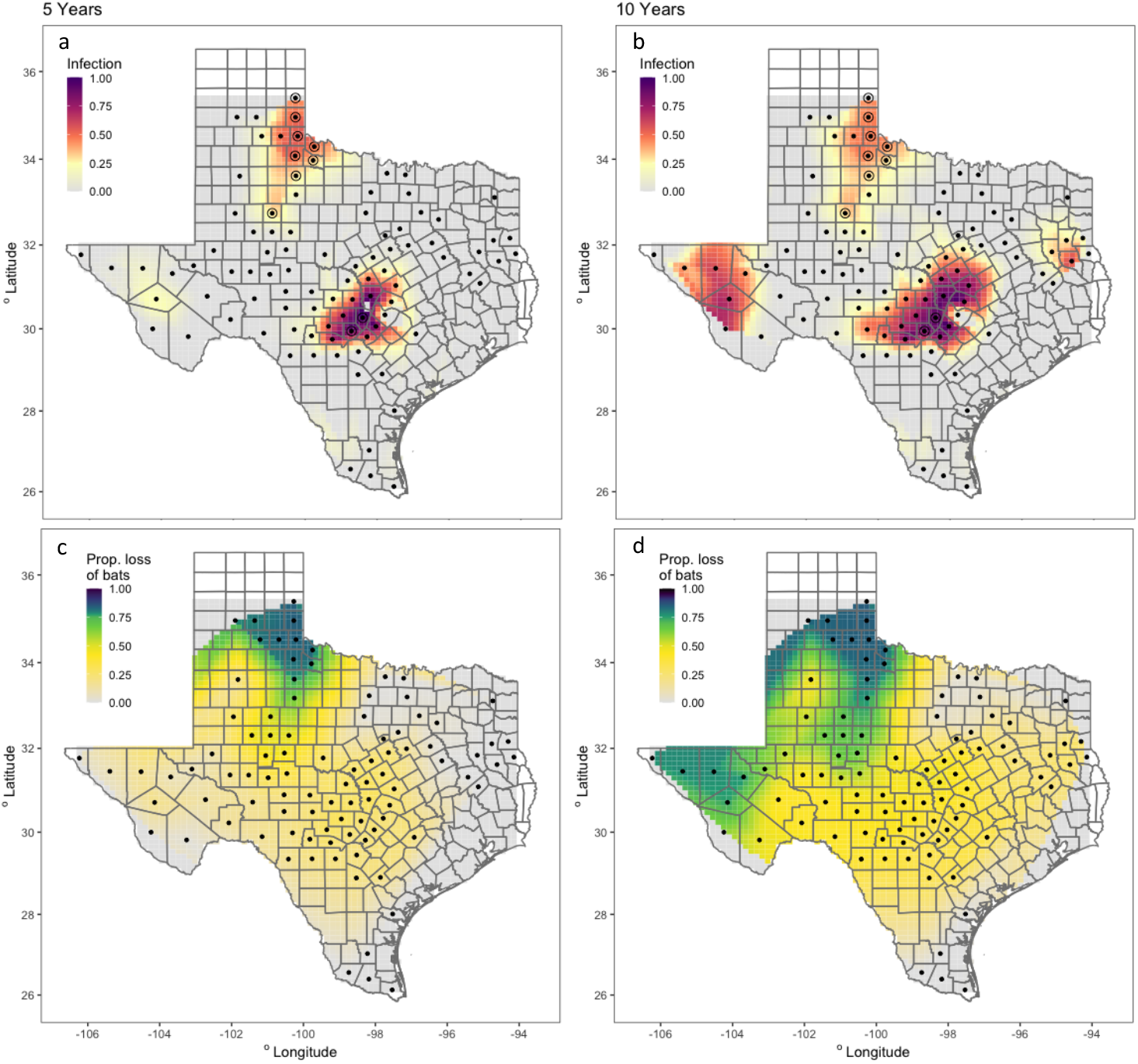
Interpolation of the carrying-capacity-scaled infection output for (**a**) 5 and (**b**) 10 years of simulation. Grey tiles show regions of 0 predicted infection. Interpolation predicts for values within a convex hull around the county centerpoints (black dots). Sites with observed infection data in 2008 marked with a circle around the point. Interpolation of the proportional loss of bats relative to infection-free model for (**c**) 5 and (**d**) 10 years. Gradient scale of heat map weighted to distinguish between larger degrees of loss (50%+).

**Figure 4.**
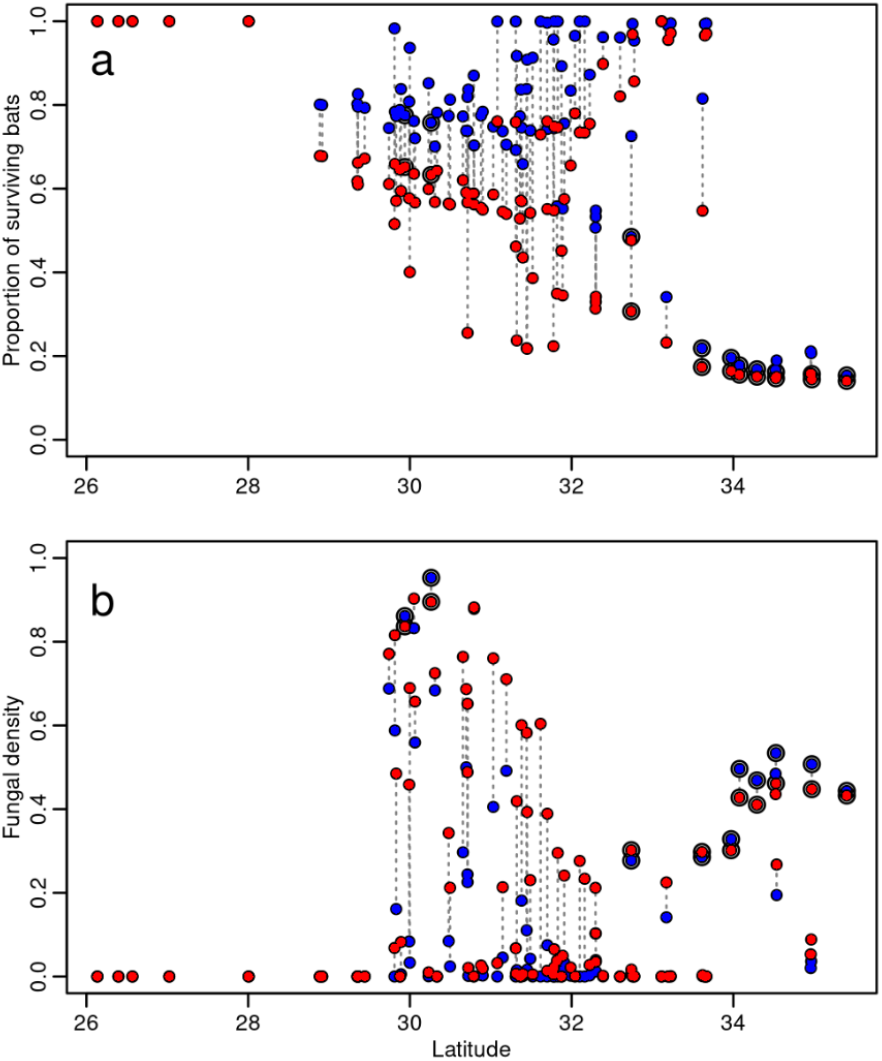
Effects of the disease on counties at different latitudes. Proportion of surviving bats (a) and the amount of free-living fungus (b), averaged for each county. The effects after 5 and 10 years are marked with blue and red dots, respectively. The matching counties are connected by dotted lines. Counties initiated with the disease are marked with a black circle around the dot. The y-axis scale on panel B is in units of fungal carrying capacity.

*Pseudogymnoascus destructans* caused low mortality in the southernmost counties under our parameter set. This is because high ambient temperatures did not support long enough hibernation periods for significant disease progression to the infectious state (Fig. 4a). The warm temperatures and resulting short hibernation period also reduced the impact of the epidemic in the central-Texas counties. While cold patches may have periods of hibernation even in warm counties, the cave temperature was then below optimal (13.0 °C, Verant et al. 2012) for fungal growth. Exposed bats carrying the fungus will be present, however, because of dispersal from affected sites.

While both transmission modes—environmental and direct—are significant components of epidemic spread, under our parameterization transmission via the environment had a larger impact, causing approximately 90% of the force of infection along the simulation time (Supplementary Material). However, sensitivity analysis on the infectivity parameters shows that similar results can be obtained by decreasing one parameter and increasing the other parameter (i.e., adjusting rate parameters associated with the two transmission modes; Supplementary Material). Removing environmental transmission from the model resulted in 99% less bat population reduction after 10 years. This occurs because most of the exposed and infected bats shed the fungus and revert back to the susceptible state during the summer, and transmissions from the environment is required to re-infect the bat population at the start of hibernation period. However, direct transmission still impacted the bat population; removing direct transmission resulted in 90% less bat population reduction. Direct transmission is the most probable cause of the disease spread into new counties. When exposed bats disperse into new sites, they shed fungus into the environment, but also transmit directly to susceptible hosts when entering hibernation. Because fungal densities remain low in the environment at new sites initially, the direct transmission route may be more prevalent.

The sensitivity analysis shows that the spread of WNS changes under variation of the parameters. Specifically, our results are most susceptible to changes in direct transmission rate, infection rate, and hibernation temperature thresholds, which increase the disease mortality and spread when the parameter value increases, and bat growth rate, which decreases mortality and spread when the parameter value increases. Increasing the mean dispersal distance (decreasing *γ*) significantly increases the number of affected patches, but does not significantly affect mortality.

## Discussion

Our predictive models found that mortality due to WNS will vary across Texas cave hibernacula, with northern sites more affected than southern sites. Results from our model suggest that in 5 to 10 years, we can expect to see a substantial (>75%) reduction in the number and size of bat populations in the northern sites because of an increase in mortality due to this fungal disease. Central sites are affected to a lesser degree, with a model projected 30–50% reduction in population densities, whereas southern sites are mostly unaffected. Interestingly, the free-living fungus reaches very high densities in central Texas where hibernation sites are most numerous, but due to warm temperatures, the bat populations in these sites are less severely affected likely due to shorter periods of torpor. The high fungal densities are in part explained by the reduced mortality, resulting in a long duration during which bats shed fungal spores.

The first documentation of WNS was anticipated to be in north Texas based on environmental characteristics (Meierhofer et al. 2019) and proximity to nearest WNS-infected sites. However, WNS was first documented on cave myotis (*Myotis velifer*) in 18 counties in central Texas in spring 2020 (TPWD 2020; Fig. 2). This is the first documentation of WNS in central and southern regions, resulting after four years of *P*. *destructans* being present in Texas. The finding of WNS in central Texas does support our model in that some regions in central Texas will have hibernacula conducive to WNS development. Our model shows that both the fungus and WNS are prevalent in central Texas, but that the proportional disease mortality is smaller in central Texas than in northern Texas. Despite the low mortality in comparison to north Texas, our model suggests that *P. destructans* reaches higher densities in central Texas than in northern Texas because fewer exposed bats succumb to the disease, increasing the continued spread of the fungus. Central Texas has the greatest abundance of known hibernacula in Texas, as well as the greatest diversity of bats in the state (Ammerman et al. 2012), increasing the potential for infection susceptibility to the disease for some bat species. Unfortunately, the origin (hibernacula) for most of the WNS positive bats is unknown, with some found—for example—outside of a house (6%), at a bridge (9%), submitted for rabies testing (31%), and on a path (3%) (J. Evans pers. com.). These WNS positive bats were found in February, suggesting these bats were recently in hibernation. However, it is still difficult to determine whether the first hibernacula with WNS are located within central Texas or elsewhere.

Our model suggests that the WNS epidemic is dependent on exposure to the spores in the environment. Indeed, contact of bats with the contaminated environment (Linder et al. 2011; Lorch et al. 2013a, 2013b) in autumn has been shown to initiate the infection (Langwig et al. 2015). Our findings further complement recent findings suggesting that persistence of high levels *P*. *destructans* in the environment result in widespread infections (Hoyt et al. 2020). Although the primary method of spread of *P*. *destructans* and WNS is bat-to-bat (Raudabaugh & Miller 2013), under our parameter set only 1 in 10 is due to direct contact to an infectious individual during hibernation. The overall pattern is not very sensitive to the relative strengths of these two components (modes of pathogen spread: environmental, direct) and temporally detailed data would be required to estimate these parameters independently. This is important to note, however, as indirect and infrequent transmission (cryptic connections) play a key role in the transmission and community-wide spread of *P. destructans* (Hoyt et al. 2018). One factor that may have resulted in the reduction in bat-to-bat transmission in our model is our choice to not include information about the Mexican free-tailed bat. Mexican free-tailed bats could greatly hasten the spread of WNS in Texas due to the proclivity to roost in large numbers, state-wide distribution, and the overlap of this species with other known WNS-affected species in cave sites (Ammerman et al. 2012). Although this species is not known to be impacted by WNS, it is a known carrier (TPWD 2018; 2019).

*Pseudogymnoascus destructans* can persist in environments in the absence of a bat host (Hoyt et al. 2015) with growth of *P. destructans* in colonies with long hibernation periods and in sites with high levels of organic detritus (Reynolds et al. 2015). Our results suggest that reducing fungal spore loads in hibernation sites may work as an effective way to slow down the epidemic spread. In north Texas where temperatures are lower than in more southern regions, bats may be experiencing longer hibernation periods than bats in southern regions. Susceptibility to the disease requires bats to stay in torpor for prolonged periods, suggesting that pathological infection occurs in regions with long periods of low ambient temperature (Ehlman et al. 2013). Indeed, knowledge of hibernation temperatures of several species in Texas (Meierhofer et al. 2019) supports the notion of extended periods of torpor of bats in north Texas than in central and southern regions of Texas. Further, the known largest bat colonies in the world exist in Texas (Ammerman et al. 2012; Schmidly & Bradley 2016), with some colonies of hibernating bat species occurring statewide in the thousands (e.g., tri-colored bat (*Perimyotis subflavus*), Sandel et al. 2001) to tens of thousands (e.g., cave myotis (*M*. velifer), Caire et al. 2019). These large colonies can provide environments conducive to the persistence of organic detritus, supporting vegetative growth of *P*. *destructans* and creating sources of increased potential environmental transmission (Reynolds et al. 2015).

Our model illustrates that infectious disease spread and infectious disease severity can become uncoupled over a gradient of environmental variation. This is particularly a product of the opportunistic nature of *P. destructans* and WNS. Whereas our model suggests some central and southern counties in Texas are less affected in terms of mortality, bats still have the potential to be exposed to *P. destructans* as some areas will have the fungus present. This is indicative of source-sink dynamics, with infected hibernacula in the north acting as sink populations and hibernacula farther south acting as source populations. The prevalence of disease as well as disease invasion rate are slow in spatially clustered landscapes (Lilley et al. 2018), as reflected in central Texas where densities of caves are greater than that of north Texas. However, movement among sites during winter (Boyles et al 2006), as well winter emergence by bats afflicted by the disease, may further the spread of WNS to uninfected areas. In southern regions of the United States such as Texas where winter activity is more common than that of northern United States (Bernard & McCracken 2017), these dynamics may be further exacerbated.

Our model-based projections are dependent on our parameterization of the dynamical model. Finding the relevant parameter set for a particular case is admittedly difficult, despite that for some parameters the values were readily available from previous work (Lilley et al. 2018). In addition, we performed an additional validation step to refine our estimates based on observations of disease spread after two years since introduction to Texas. Our initial best-estimate parameter values undershot the observed pattern of disease prevalence in 2020, and the refined estimates after validation led to faster spread and less lethal disease (Supporting Information). The values we chose for rejection thresholds (rejection criteria) also reflect our subjective views of at which level of prevalence the WNS disease and *P. destructans* would be detected with the effort the surveys were performed. Similar analysis and model projections could be performed with our model framework in the following years when new data become available, thereby refining and improving estimates and predictions.

We anticipate that the spread of *P*. *destructans* will be slow and display source-sink dynamics in Texas and at more southern regions such as northern Mexico. We further anticipate that the spread of *P. destructans* in central Texas—where caves are more clustered—will be similar to eastern United States, where the spread increased with proximity to nearest infected site (Ingersoll et al. 2016). Indeed, results from our model suggest that conservation actions should consider sites that have temperatures conducive to hibernation and suitable for fungal growth, as these sites may be more susceptible to local extinctions given source-sink dynamics. In light of these factors, and that large colonies of bats in temperatures suitable for *P*. *destructans* experience the greatest impacts of WNS (Wilder et al. 2011), we recommend prioritizing the preservation of large overwintering colonies of bats in north Texas through management actions. We further propose that future work should focus on within-season movement movements of cavernicolous bat species forming large colonies at southern latitudes to ascertain infectiousness, and thus the rate of spread amongst these populations.

## Supporting information

Hibernaculum Temperature Modeling

Model description and additional analyses

## Supporting Information

Model description and additional analyses (Appendix S1), Hibernaculum temperature modelling (Appendix S2) are available online. The authors are solely responsible for the content and functionality of these materials. Queries (other than the absence of the material) should be directed to the corresponding author.

## Acknowledgements

D. Wright, Dr. Comer, Dr. Godwin are thanked for site access; J. Kennedy is thanked for site access and data collection; K. Demere, B. Stamps, L. Wolf, S. Goree, J. Carey are thanked for field assistance in data collection. Funding for this project was provided through the U.S. Fish and Wildlife Service’s State Wildlife Grant Program (CFDA# 15.611) as administered by Texas Parks and Wildlife Department and the U.S. Fish and Wildlife Service’s (Service) CFDA Program (15.657). Additional funding was provided by the fightWNS ‘Micro Grants for Microbats’ and the NSS WNS Rapid Response Fund. This work was supported by the Fulbright Finland Foundation and Finnish National Agency for Education EDUFI (MBM) and the Academy of Finland (TML, grant# 331515).

