## Supplementary material for "Ten-year projection of white-nose syndrome disease dynamics at the southern leading-edge of infection in North America": Hibernaculum Temperature Modeling: Supplement.html

SUPPLEMENT: Cave Temperature Estimation


Code 

- Show All Code
- Hide All Code
- Download Rmd

### SUPPLEMENT: *Cave Temperature Estimation*

### 1 Intro

Within-cave conditions are an essential part of our model, but sufficient data for modeling these conditions is not available. To overcome this limitations, we try to model the conditions within caves, based on the ambient conditions outside the caves. We also account for cave location, as the association between above and below ground temperature conditions might vary across sites.

### 2 Data

We use both measurements from within a collection of caves, as well as Texas-wide PRISM data.

#### 2.1 PRISM data


```
library(data.table)
library(lubridate)
library(dplyr)

ambient = fread('Weather Data/PRISMAllCounties.csv',sep=',') %>%
  select(County = Name, everything()) %>%
  mutate(Date = mdy(Date),
         TempRNG = TempMaxC - TempMinC)
ambient[sample(nrow(ambient),10),]
```

#### 2.2 Cave data


```
caves = fread('All cave data.txt',sep = '\t')
caves$Date = as.Date(strptime(caves$Date,format = '%m/%d/%y'))
names(caves) = gsub(' ','_',names(caves))

caves %>% mutate(month = month(Date)) %>% 
  group_by(County, month) %>% 
  summarise(`Mean Temperature` = mean(Celsius),
            `Mean Humidity` = mean(Humidity),
            `Mean dew point` = mean(Dew_Point)) %>%
  mutate_if(is.numeric, round, 2) %>%
  filter(month %in% c(10:12,1,2))
```

#### 2.3 Data manipulation

Calculate aggregate values from data mmeasured within-caves:


```
caves %>%
  group_by(Site, Longitude, Latitude, 
           County,Date, Winter_Bats) %>%
  summarize(muT = mean(Celsius),
            minT = min(Celsius),
            maxT = max(Celsius),
            sdT = sd(Celsius)) %>%
  select(Site_lat = Latitude, Site_lon = Longitude,
         everything()) -> avgs

avgs %>% ungroup() %>% slice(sample(n(),10)) %>% select(-contains('Site'))
```


Combining the the ambient date with cave aggregates:


```
full = inner_join(avgs,ambient) %>%
  mutate(month = month(Date))

full %>% ungroup() %>% slice(sample(n(),10)) %>% select(-contains('Site'))
```


Filter data to include only winter months (November-February), and exclude records with average within-cave temperature above 20 degrees (also exclude two sites with bad-quality data):


```
df_tr = full %>% 
  filter(month%in%c(10:12,1:2) & muT < 20,
         Site!='Deep' & Site!='Cave Spring')

df_tr %>% ungroup() %>% 
  slice(sample(n(),10)) %>% select(-contains('Site'))
```


Visualize association between ambient and within-cave temperature:


```
library(ggplot2)
library(tidyr)
df_tr %>% select(muT, minT, maxT, sdT, TempMeanC) %>% ungroup() %>%
  gather(key='Variable', value='value',
         -County, -Site, -Site_lon, -Site_lat, 
         -Date, -starts_with('Temp')) %>%
ggplot(aes(TempMeanC, value)) +
  geom_point(shape=21, alpha=0.5, fill='gray50') +
  facet_wrap(~Variable, ncol=2, scales = 'free_y') +
  theme_bw() +
  theme(legend.position = 'n') +
  geom_smooth() +
  xlab('Mean Ambient Temperature (°C)') +
  ylab('Within-cave Temperature (°C)')
```


There is a clear association between ambient and within-cave average temprerature. While this is expected, the association is not quite linear, also because we restricted the cave data to top at 20°C.

While the within-cave temperarure variability (SD) seems to be rather independent of ambient conditions, the minimum and maximum are associated with outside-cave conditions.

### 3 Modeling

Given the observed non-linear association between ambient and within-cave conditions, we considered using GAM-models to best capture the relationship. However, we had to abandon this approach, due to overfitting problems. In the end, we wanted to be conservative with our estimates and therefor used simple linear models.

We will train a model of mean temperature as well as min/max temperature. The models use month, a second-order polynomial of the mean ambient temperature, as well as the site location as independent variables. To get a more reliable idea of the generality of the model, we use 10-fold cross validation to get the coefficient of determination:


```
library(caret)
trc = trainControl(method='cv', number=10)
formulas = lapply(c('muT','minT','maxT'), function(x){
  formula(paste(x,'~month+TempMeanC+I(TempMeanC^2)+Site_lat+Site_lon'))
})
models = lapply(formulas, function(x){
  train(x,data = df_tr, method='lm', trControl = trc)
})
names(models) = c("mean", "min", "max")
```


```
lapply(models, summary)
```


```
$mean

Call:
lm(formula = .outcome ~ ., data = dat)

Residuals:
    Min      1Q  Median      3Q     Max 
-7.5895 -0.9725 -0.0344  0.8811 10.2775 

Coefficients:
                  Estimate Std. Error t value Pr(>|t|)    
(Intercept)      63.652392   3.723868  17.093  < 2e-16 ***
month             0.536641   0.014076  38.126  < 2e-16 ***
TempMeanC         0.322524   0.029173  11.055  < 2e-16 ***
`I(TempMeanC^2)` -0.004501   0.001198  -3.755 0.000185 ***
Site_lat         -0.844177   0.041061 -20.559  < 2e-16 ***
Site_lon          0.289714   0.038537   7.518 1.41e-13 ***
---
Signif. codes:  0 ‘***’ 0.001 ‘**’ 0.01 ‘*’ 0.05 ‘.’ 0.1 ‘ ’ 1

Residual standard error: 1.943 on 851 degrees of freedom
Multiple R-squared:  0.7695,    Adjusted R-squared:  0.7682 
F-statistic: 568.2 on 5 and 851 DF,  p-value: < 2.2e-16


$min

Call:
lm(formula = .outcome ~ ., data = dat)

Residuals:
     Min       1Q   Median       3Q      Max 
-10.0479  -0.9672   0.0637   1.1248   8.7513 

Coefficients:
                  Estimate Std. Error t value Pr(>|t|)    
(Intercept)      59.934645   4.122225  14.539  < 2e-16 ***
month             0.615625   0.015581  39.511  < 2e-16 ***
TempMeanC         0.337795   0.032294  10.460  < 2e-16 ***
`I(TempMeanC^2)` -0.004339   0.001327  -3.271  0.00112 ** 
Site_lat         -0.874839   0.045453 -19.247  < 2e-16 ***
Site_lon          0.256205   0.042660   6.006 2.82e-09 ***
---
Signif. codes:  0 ‘***’ 0.001 ‘**’ 0.01 ‘*’ 0.05 ‘.’ 0.1 ‘ ’ 1

Residual standard error: 2.151 on 851 degrees of freedom
Multiple R-squared:  0.7668,    Adjusted R-squared:  0.7655 
F-statistic: 559.7 on 5 and 851 DF,  p-value: < 2.2e-16


$max

Call:
lm(formula = .outcome ~ ., data = dat)

Residuals:
    Min      1Q  Median      3Q     Max 
-6.5249 -1.0297 -0.1624  0.7879 14.4530 

Coefficients:
                  Estimate Std. Error t value Pr(>|t|)    
(Intercept)      66.656378   4.083319  16.324  < 2e-16 ***
month             0.452404   0.015434  29.312  < 2e-16 ***
TempMeanC         0.323283   0.031989  10.106  < 2e-16 ***
`I(TempMeanC^2)` -0.005142   0.001314  -3.913 9.86e-05 ***
Site_lat         -0.807511   0.045024 -17.935  < 2e-16 ***
Site_lon          0.318644   0.042257   7.541 1.20e-13 ***
---
Signif. codes:  0 ‘***’ 0.001 ‘**’ 0.01 ‘*’ 0.05 ‘.’ 0.1 ‘ ’ 1

Residual standard error: 2.131 on 851 degrees of freedom
Multiple R-squared:  0.6946,    Adjusted R-squared:  0.6928 
F-statistic:   387 on 5 and 851 DF,  p-value: < 2.2e-16
```


The model seems to capture the variation in within-cave temperature mean, with an adjusted *R*-squared of 0.77. The temperature extremes have similar predictability.

Next, we calculate model predictions:


```
df_tr %>% select(County, Date, 
                 mean_obs = muT, 
                 min_obs = minT,
                 max_obs = maxT) %>% ungroup() -> x
predicted = sapply(models, function(x) predict(x, df_tr))
df_pred = cbind(x, predicted)

df_pred %>% ungroup() %>% mutate_if(is.numeric, round, 2) %>%
  slice(sample(n(),10)) %>% select(-contains('Site'))
```


Visualize predictions VS. observed values:


```
select(df_pred, ends_with('obs')) %>% 
  gather(key='variable',value='observed') %>% 
  mutate(variable = gsub('_obs', '', variable)) %>%
  bind_cols(
    select(df_pred, mean, min, max) %>% 
      gather(key='variable',value='predicted') %>%
      select(-variable)
  ) %>%
ggplot(aes(observed,predicted)) +
  geom_point(col='blue3',alpha=0.5, size=1) + 
  geom_abline(intercept = 0,slope = 1, 
              col = 'red3',size=1,alpha=.5) +
  facet_wrap(~variable, ncol=3) +
  theme_minimal() +
  xlab('True Temperature (°C)') +
  ylab('Estimated Temperature (°C)')
```

### 4 Aggregate estimates


```
ambient %>% 
  select(County, TempMeanC, Date,
         Site_lat = Latitude, 
         Site_lon = Longitude) -> df_tp
df_tp$month = month(df_tp$Date)
df_tp %>% filter(month %in% c(1,2,10:12)) -> df_tp
df_tp %>%  
bind_cols(
    as.data.frame(
      sapply(models, function(x) predict(x, df_tp))
      )
  ) -> df_tp
```


Next, we calculate estimates for mean/min/max of within-cave temperature. All are calculated at the level of Texas, as well as individual counties:


```
df_tp %>% ungroup() %>% 
  mutate(overall_mean = mean(mean),
         overall_min = min(min),
         overall_max = max(max)) %>%
  mutate_if(is.numeric,round, 2) %>%
  group_by(County, overall_mean, overall_min, overall_max) %>%
  summarize(mean_T = mean(mean),
            min_T = min(min),
            max_T = max(max)) %>%
  mutate_all(round, 2) -> out
out
```


The mean for each county will be used as the mean cave temperature in the dynamical model. The variation around this mean, however, is taken as the variance of model residuals:


```
round(var(resid(models$mean)),2)
```


```
[1] 3.75
```


The minima and maxima are used for setting ranges for dynamic temperature varion within caves in the model.
