## Supplementary material for "Ten-year projection of white-nose syndrome disease dynamics at the southern leading-edge of infection in North America": Model description and additional analyses

#### S1.1. Model

The model, modified from Lilley et al. 2018, is described by the following system of differential equations:

| | | |
|---|---|---|
| | $\frac{d}{dt}s^a_{i,j} = r_h(T^o_j, T^c_{i,j})(s^a_{i,j} + e^a_{i,j}) - r_h(T^o_j, T^c_{i,j})s^a_{i,j}\frac{N_{i,j}}{K_{i,j}} + \delta_e e^a_{i,j} - \frac{\beta_e}{K_{i,j}}s^a_{i,j}g(f_{i,j}) + \omega_{a\leftarrow h}(T^o_j, T^c_{i,j})s^h_{i,j} - \omega_{h\leftarrow a}(T^o_j, T^c_{i,j})s^a_{i,j}$ | (S1.1) |
| | $\frac{d}{dt}e^a_{i,j} = -r_h(T^o_j, T^c_{i,j})e^a_{i,j}\frac{N_{i,j}}{K_{i,j}} + \delta_e e^a_{i,j} + \frac{\beta_e}{K_{i,j}}s^a_{i,j}g(f_{i,j}) + \delta_n n^a_{i,j} - \delta_e e^a_{i,j} + \omega_{a\leftarrow h}(T^o_j, T^c_{i,j})e^h_{i,j} - \omega_{h\leftarrow a}(T^o_j, T^c_{i,j})e^a_{i,j}$ | (S1.2) |
| | $\frac{d}{dt}n^a_{i,j} = -r_h(T^o_j, T^c_{i,j})n^a_{i,j}\frac{N_{i,j}}{K_{i,j}} - \delta_n n^a_{i,j} + \omega_{a\leftarrow h}(T^o_j, T^c_{i,j})n^h_{i,j} - \omega_{h\leftarrow a}(T^o_j, T^c_{i,j})n^a_{i,j}$ | (S1.3) |
| | $\frac{d}{dt}s^h_{i,j} = -\frac{\beta_d}{K_{i,j}}s^h_{i,j}n^h_{i,j} - \mu_h s^h_{i,j} + \omega_{h\leftarrow a}(T^o_j, T^c_{i,j})s^a_{i,j} - \omega_{a\leftarrow h}(T^o_j, T^c_{i,j})s^h_{i,j}$ | (S1.4) |
| | $\frac{d}{dt}e^h_{i,j} = \frac{\beta_d}{K_{i,j}}s^h_{i,j}n^h_{i,j} - \mu_h e^h_{i,j} - \varphi e^h_{i,j} + \omega_{h\leftarrow a}(T^o_j, T^c_{i,j})e^a_{i,j} - \omega_{a\leftarrow h}(T^o_j, T^c_{i,j})e^h_{i,j}$ | (S1.5) |
| | $\frac{d}{dt}n^h_{i,j} = \varphi e^h_{i,j} - \mu_h n^h_{i,j} - \mu_f n^h_{i,j} + \omega_{h\leftarrow a}(T^o_j, T^c_{i,j})n^a_{i,j} - \omega_{a\leftarrow h}(T^o_j, T^c_{i,j})n^h_{i,j}$ | (S1.6) |
| | $\frac{d}{dt}f_{i,j} = r_f f_{i,j} h(T^c_{i,j}) - \frac{r_f}{K_{i,j}}f_{i,j}f_{i,j} + \lambda(n^a_{i,j} + n^h_{i,j})$ | (S1.7) |

The subindices *i* and *j* denote patches and counties, and superindices *a* and *h* denote hosts in active and hibernating states, respectively. The system state is defined by susceptible, exposed, and infectious bats, denoted by *s*, *e*, and *n*, in both active and hibernating states, and additionally the free-living fungus, denoted by *f*. The sum of active bats is denoted by $N_{i,j} = s^a_{i,j} + e^a_{i,j} + n^a_{i,j}$. Parameters are listed in main text (ref).

Patches and counties vary in ambient temperatures $T^o_j$, which has the same value for all patches in a county, and hibernaculum temperatures $T^c_{i,j}$, to which patches within a county are binned following a Gaussian distribution (see main text). Both vary sinusoidally along a year. The ambient temperatures are calculated as:

| | | |
|---|---|---|
| | $$T_j^o(t) = a_j^o \sin\frac{2\pi t}{365} + b_j^o$$ | (S2) |

Similarly, hibernaculum temperatures are calculated as:

| | | |
|---|---|---|
| | $$T_{i,j}^c = a_{i,j}^c \sin\frac{2\pi t}{365} + b_{i,j}^c$$ | (S3) |

Transition rates between active and hibernation states, and bat growth and density dependence, are switched on and off depending on whether a patch is in hibernation or not. In addition to transitions and bat growth, hibernation affects dispersal. Dispersal between two patches occurs only when both patches are in active state. A patch is considered to be in hibernation ($\pi = 1$) when either ambient temperature is below a threshold value $\eta_{h\leftarrow a,1}$ or ambient temperature is below a higher threshold value $\eta_{h\leftarrow a,2}$ and cave temperature is below $\eta_{h\leftarrow a,1}$. Conversely, a patch is not in hibernation ($\pi = 0$) when these requirements are not met. Formally, this is defined as:

| | | |
|---|---|---|
| | $$\pi(T_j^o, T_{i,j}^c) = \begin{cases} 1 \; if \; T_j^o \le \eta_{h\leftarrow a,1}^o \\ 1 \; if \; T_j^o \le \eta_{h\leftarrow a,2}^o \; and \; T_{i,j}^c \le \eta_{h\leftarrow a}^c \\ 0 \; otherwise \end{cases}$$ | (S4) |

Transition rates and bat growth are then defined in terms of the hibernation state $\pi$ in a patch. Transition into hibernation is:

| | | |
|---|---|---|
| | $$\omega_{h\leftarrow a}(T_j^o, T_{i,j}^c) = \pi(T_j^o, T_{i,j}^c) \cdot \widehat{\omega}_{h\leftarrow a}$$ | (S5) |

and transition into active state is:

| | | |
|---|---|---|
| | $$\omega_{a\leftarrow h}(T_j^o, T_{i,j}^c) = \left(1 - \pi(T_j^o, T_{i,j}^c)\right) \cdot \widehat{\omega}_{a\leftarrow h}$$ | (S6) |

Bat growth is defined as:

| | | |
|---|---|---|
| | $$r_h(T_j^o, T_{i,j}^c) = \left(1 - \pi(T_j^o, T_{i,j}^c)\right) \cdot \hat{r}_h$$ | (S7) |

Note, that hibernation transitions follow simple step functions, whereas Lilley et al. 2018 used smooth sigmoidal transitions between active and hibernating states. We have chosen these forms in order to achieve simpler parameterisation and to account for the combined effect of ambient and hibernaculum temperatures.

Growth of the free-living fungus is affected by the cave temperature $T_{i,j}^c$ as:

| | | |
|---|---|---|
| | $$h(T_{i,j}^c) = \begin{cases} 0 \; if \; T_{i,j}^c \le T_{min}^c \\ h_{max}\left(c_1(T_{i,j}^c - T_{min}^c)\right)^2 \left(1 - e^{c_2(T_{i,j}^c - T_{max}^c)}\right) \; if \; T_{min}^c < T_{i,j}^c < T_{max}^c \\ 0 \; if \; T_{i,j}^c \ge T_{max}^c \end{cases}$$ | (S8) |

Similarly to Lilley et al. 2018, we used the parameters $h_{max} = 50$, $T_{min}^c = 7$, $T_{max}^c = 15$, $c_1$ = 0.0377, $c_2$ = 0.25 for the fungal growth function, resulting in an optimum at around 13°C.

The shape of force of infection for environmental transmission depends on the function $g(f_{i,j})$. Lilley et al. 2018 assumed a sigmoidal shape because of theoretical considerations (ref Anttila). Here, we assume a simpler linear shape, i.e. $g(f_{i,j}) = f_{i,j}$ for the main simulations and results communicated in main text. In addition, we have considered the case where the force of infection assumes a sigmoidal shape:

| | | |
|---|---|---|
| | $$g(f_{i,j}) = \beta_{e,max} \frac{\left(\frac{f_{i,j}}{ID_{50}}\right)^k}{1 + \left(\frac{f_{i,j}}{ID_{50}}\right)^k}$$ | (S9) |

We report results below in section S3.

### S1.2. Sensitivity analysis

We analyzed the sensitivity of simulated outcomes to variations in model parameters. The starting point for each parameter was our best-guess value (Table T, main text), which was varied up and down by 5% and 10% increments. For each variation we calculated the relative change in a) overall mortality (change in live bat population density compared to infection-free scenario, in patch units) and b) number of affected counties (counties with more than 5% reduction in bat population compared to disease-free scenario), 5 and 10 years after introduction of the *pd* fungus. The results are shown in figures S1-S4.

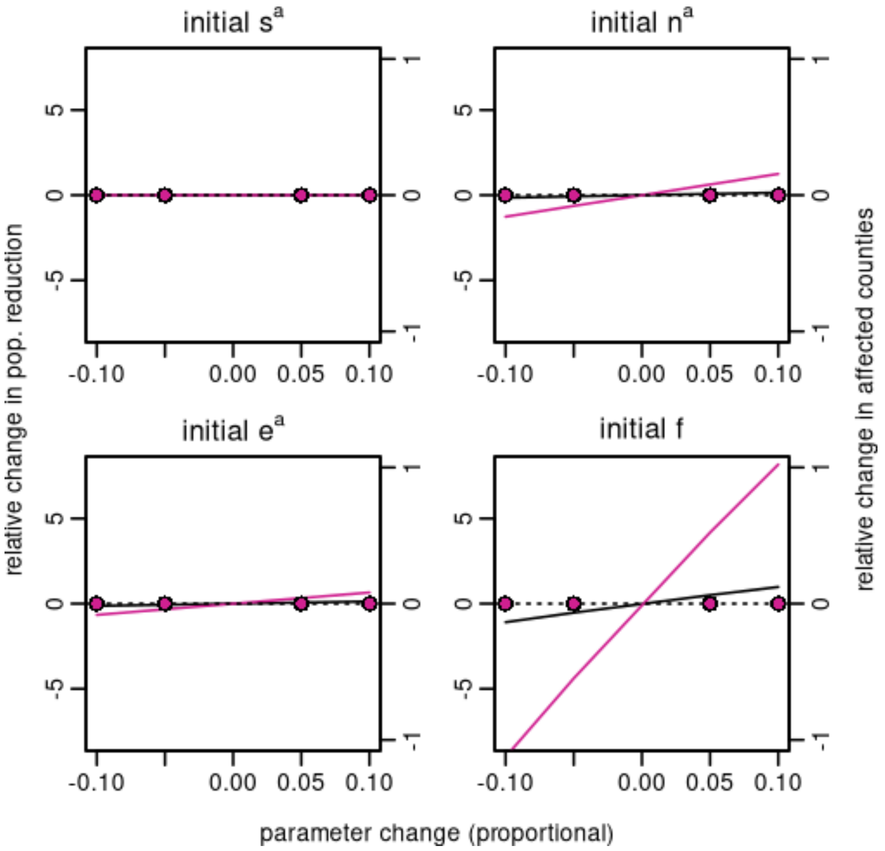


**Appendix S1**. Sensitivity of simulated outcomes to initial conditions in initially affected patches (2017). Lines show difference in overall mortality (population density change compared to disease-free scenario), and dots show difference in the number of affected counties (> 5% reduction in bat population compared to disease-free scenario). The difference after 5 years is shown in purple and the difference after 10 years in black. Note that the y-axis units are different for lines and dots. Overall mortality is expressed in patch / $K_{ij}$ units (approx. 900 bats), and affected counties may have varying numbers of patches. The horizontal dotted line of no effect is shown for comparison.

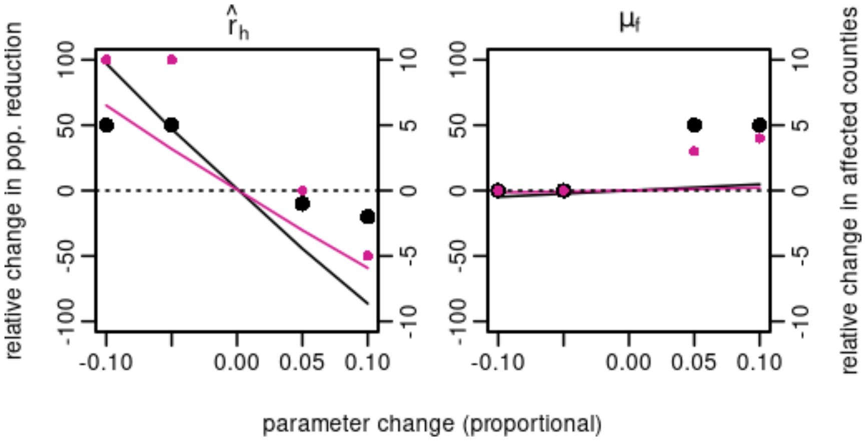


**Appendix S2**. Sensitivity of simulated outcomes to bat population growth and hibernation mortality parameters. See Figure S1 caption for full explanation.

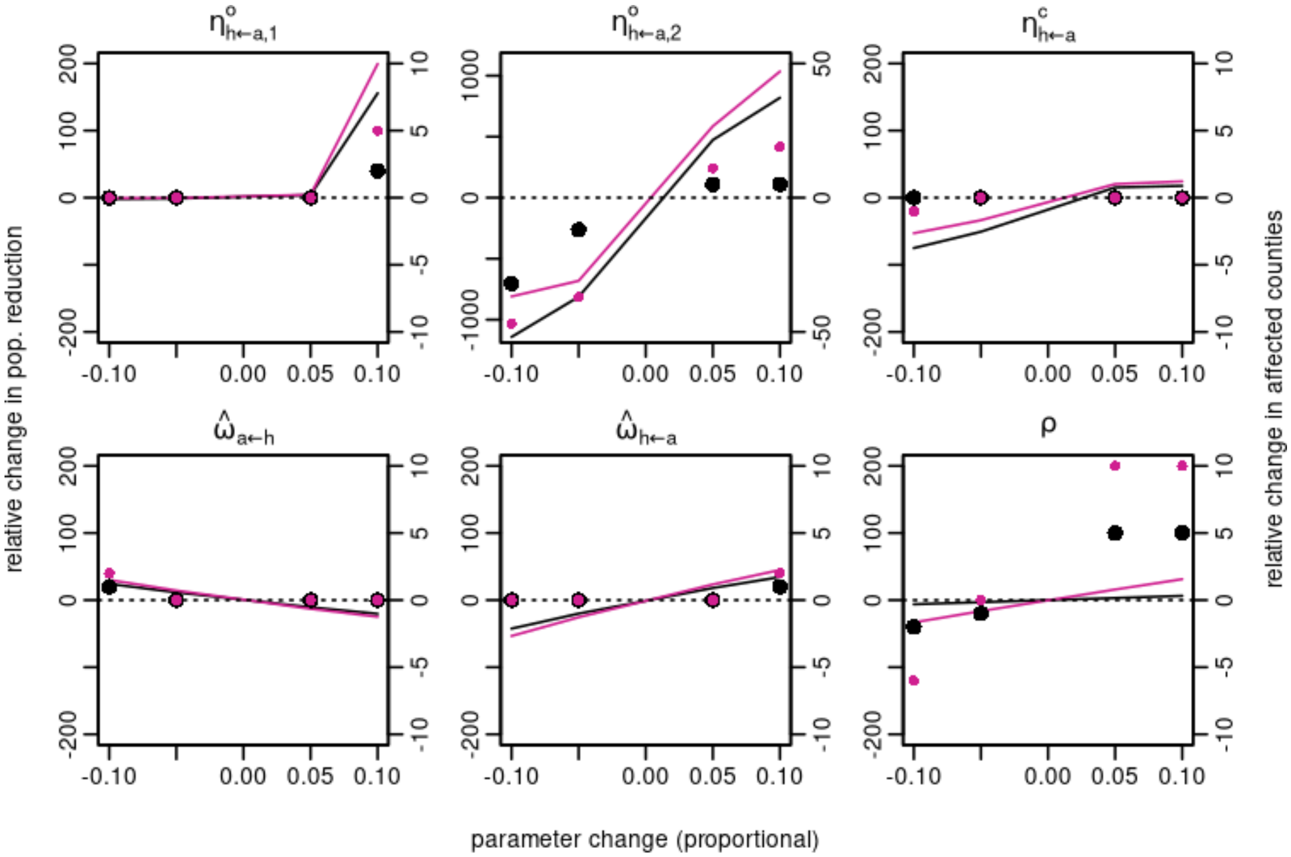


**Appendix S3**. Sensitivity of simulated outcomes to hibernation parameters and migration proportion. See Figure S1 caption for full explanation.

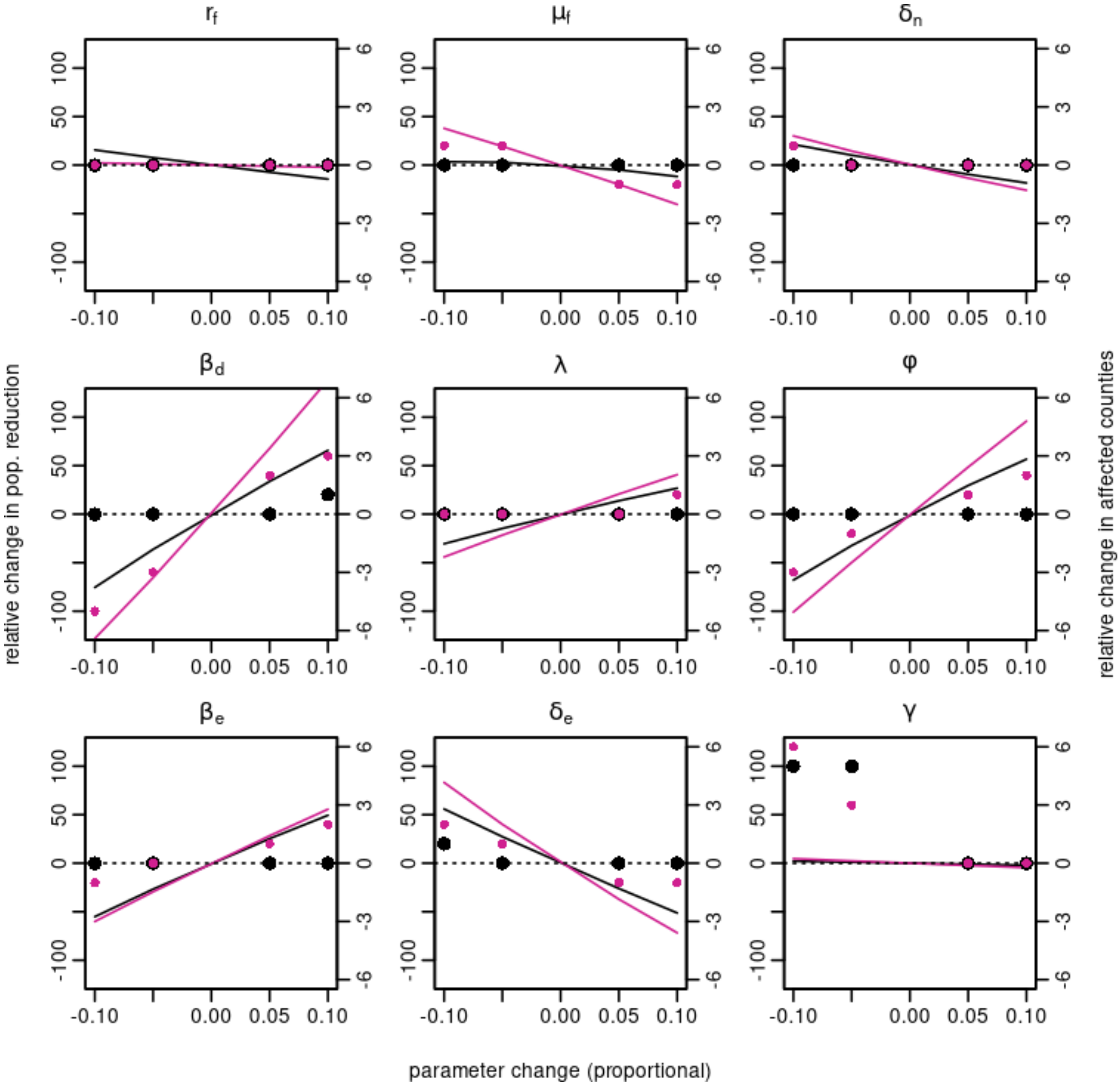


**Appendix S4**. Sensitivity of simulated outcomes to various model parameters. See Figure S1 caption for full explanation

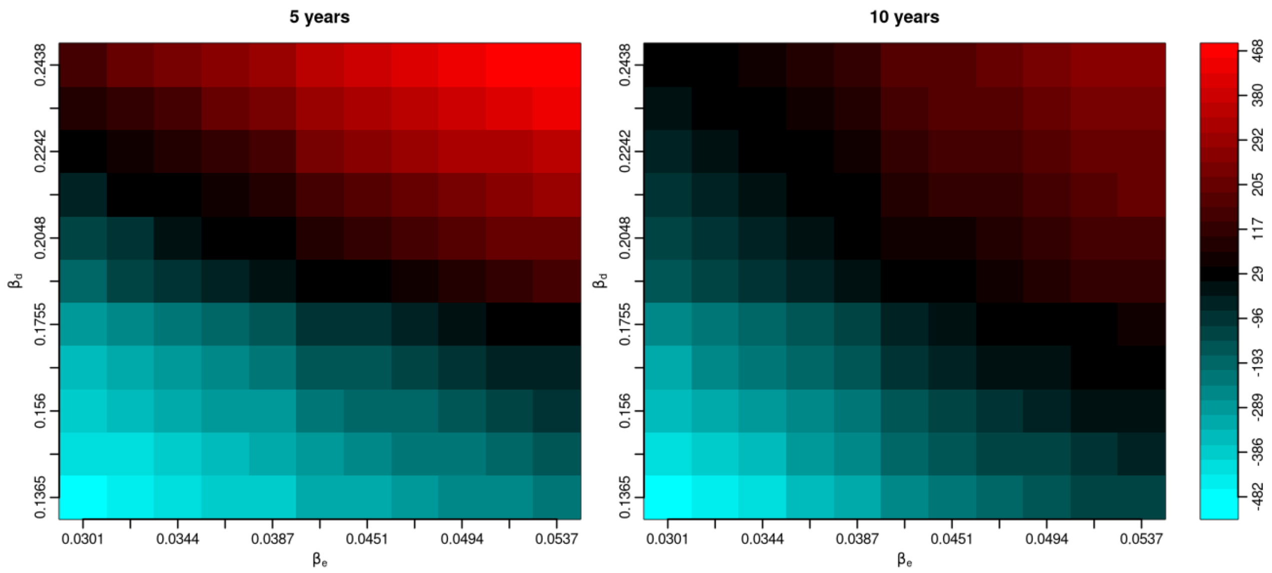


**Appendix S5**. Sensitivity of simulated population reduction to direct and environmental infectivity components. Red areas show higher reduction in bat population due to the fungal disease and blue areas show less reduction with respect to the base parameter set (center on *x* and *y* axes). Population reduction is expressed in units of *K*, *i.e.* hibernation site carrying capacities.

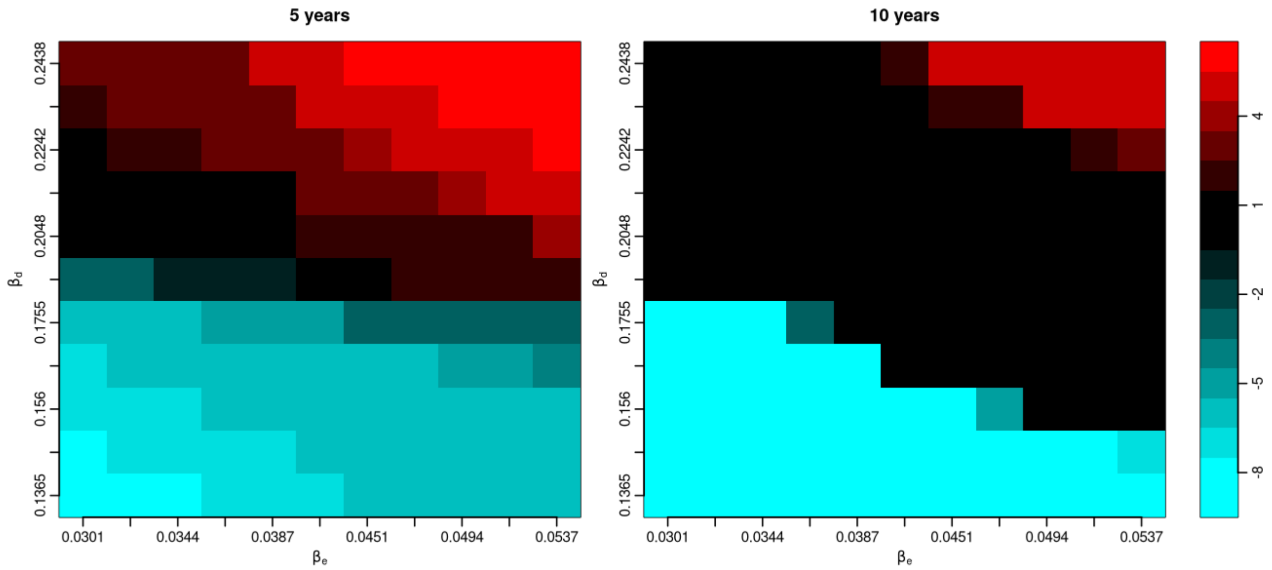


**Appendix S6**. Sensitivity of affected patches to direct and environmental infectivity components. Red areas show increase in the number of affected patches, and blue areas show decrease with respect to the base parameter set (center on *x* and *y* axes).

### S1.3. Sigmoidal force of infection

In addition to using a linear force of infection as a function of pathogen density for environmental transmission, we tested a sigmoidal response (equation S9). The sigmoid curve can be parameterized in various ways compared to the simple linear form. For simplicity, we decided to fix $\beta_{e,max} = \beta_e$ and $k = 4$ and vary $ID_{50}$ to get a range of different curves. The sigmoid function begins with an overproportional (convex) part, which in our case is below the linear form, and then crosses the linear function to an underproportional (concave) part, which may reach above the linear form. For a desired crossing point $z$ we can find $ID_{50}$ following:

$$ID_{50} = \left(z^k \frac{1-z}{z}\right)^{\frac{1}{k}} \quad \text{(S10)}$$

We explore the outcomes under sigmoidal infectivity functions with crossing points at 0.1 – 0.7. Results are shown in Figure S5.

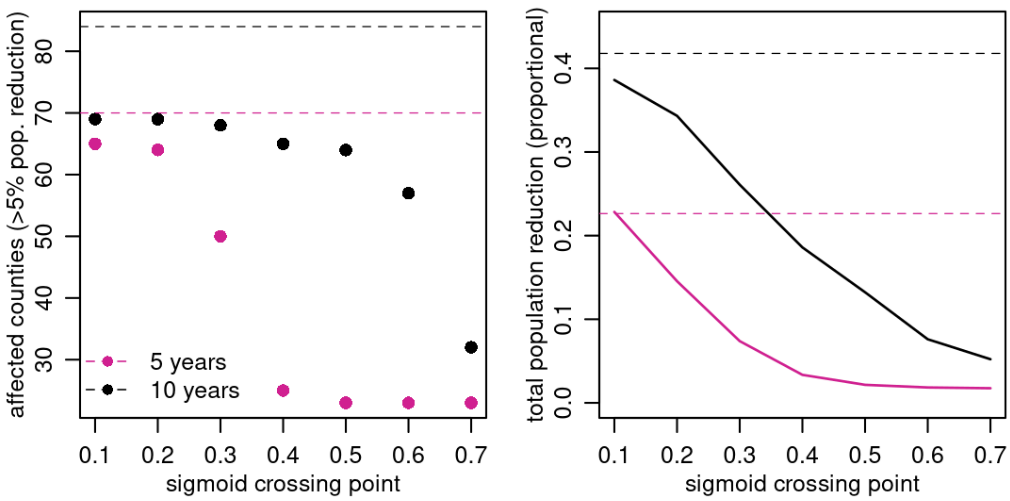


**Appendix S7**. Simulated outcomes under sigmoidal force of infection for environmental transmission. The left panel shows the difference in the number of affected counties (> 5% reduction in bat population compared to disease-free scenario). The right panel shows the difference in overall mortality (population density change compared to disease-free scenario). The difference after 5 years is shown in purple and the difference after 10 years in black. Overall mortality is expressed in patch / $K_{ij}$ units (approx. 900 bats), and affected counties may have varying numbers of patches. The horizontal dashed lines show corresponding values under linear force of infection for comparison.

### S1.5. Simulations without environmental or direct transmission components

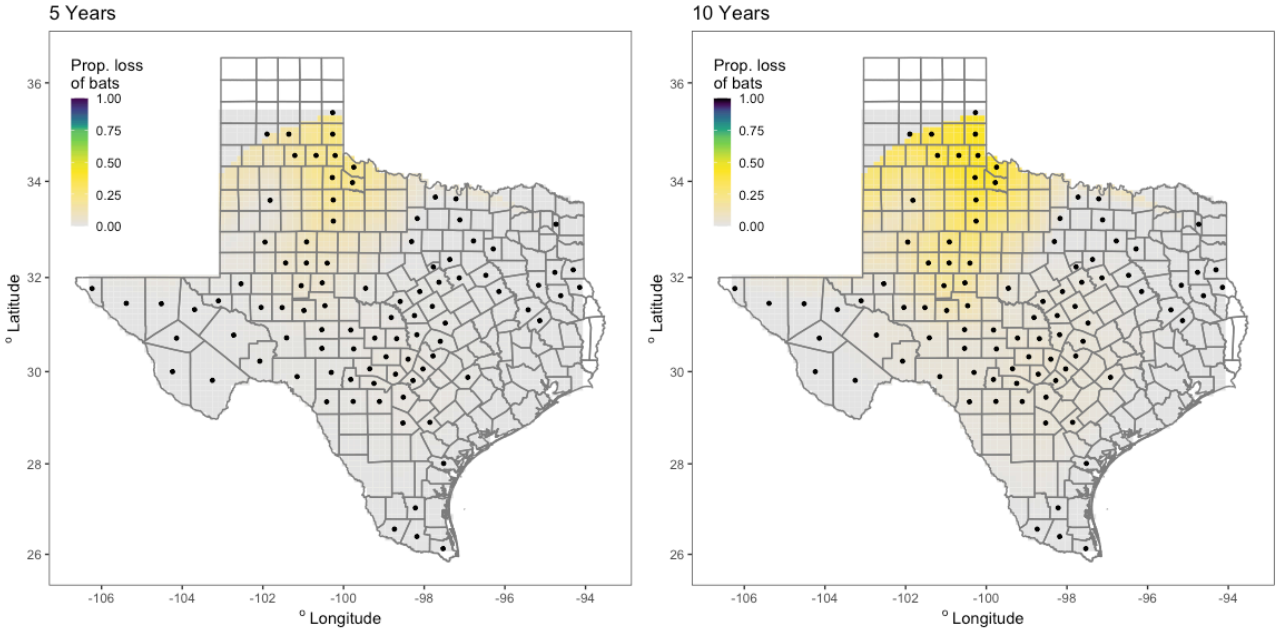


**Appendix S8.** Interpolation of the proportional loss of bats under environmental transmission only (no direct transmission component) relative to infection-free model for (**a**) 5 and (**b**) 10 years.

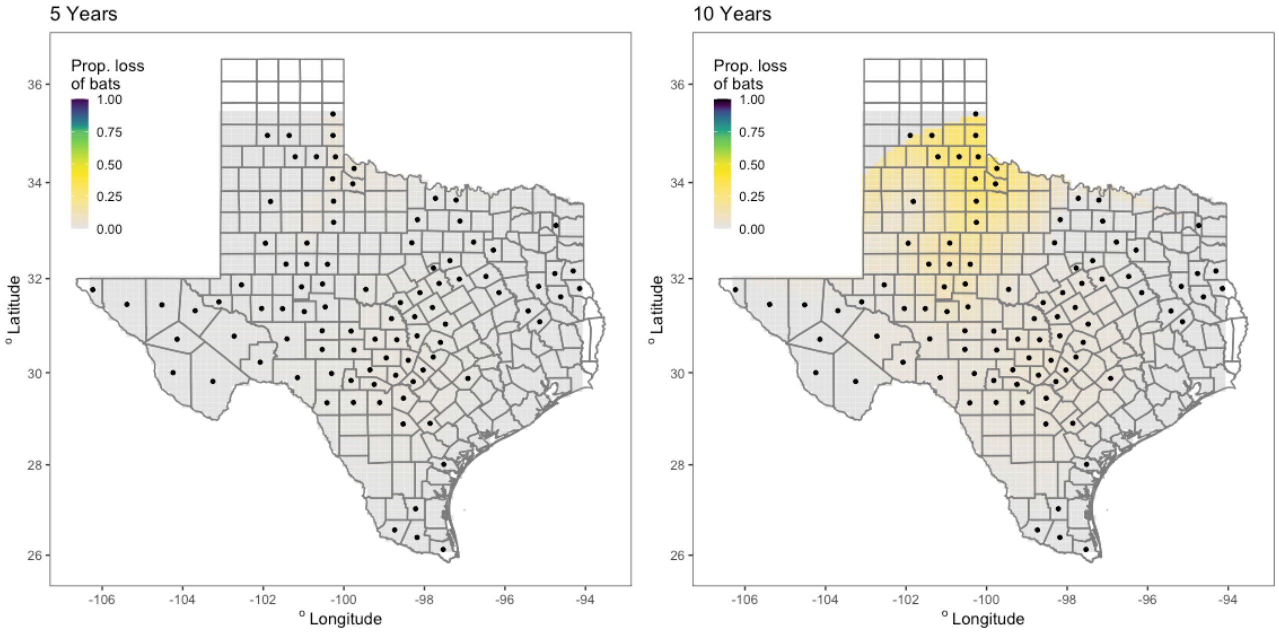


**Appendix S9.** Interpolation of the proportional loss of bats under direct transmission only (no environmental transmission component) relative to infection-free model for (**a**) 5 and (**b**) 10 years.

### S1.6. Total population dynamics

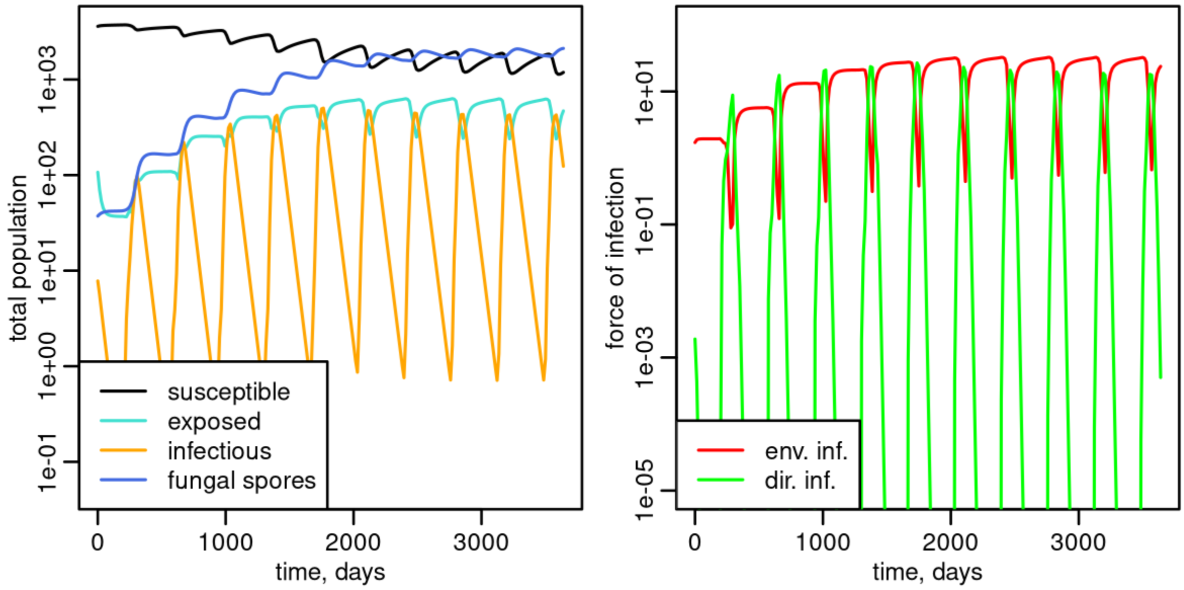


**Appendix S10.** Total population dynamics (left panel) and force of infection (right panel) under our validated parameter set for the full simulation period of 10 years.

### S1.7. Parameter validation

**Appendix S11.** The initial estimated parameter values and our estimates after the validation step based on 2020 WNS survey data (see Materials and Methods for details). Our prior parameters had each parameter independently distributed according to Galton (i.e. log-Gaussian) distribution with exp(μ) = the value listed below (Initial estimate) and exp($\sigma^2$) = 1.

| Symbol | Parameter name | Initial estimation | Value after validation |
|---|---|---|---|
| $\hat{r}_h$ | Bat population growth rate | 0.00333 | 0.00333 |
| $r_f$ | Fungal growth rate | 0.00143 | 0.00152 |
| $\beta_e$ | Environmental transmission rate | 0.0343 | 0.043 |
| $\beta_d$ | Direct transmission rate | 0.0983 | 0.195 |
| $\mu_h$ | Hibernation mortality | 0.00117 | 0.0012 |
| $\mu_f$ | Disease mortality | 0.0662 | 0.039 |
| $\lambda$ | Fungal shedding | 0.0117 | 0.017 |
| $\delta_e$ | Recovery (exposed to susceptible) | 0.0933 | 0.0488 |
| $\delta_n$ | Recovery (infectious to exposed) | 0.0290 | 0.0225 |
| $\varphi$ | Infection rate | 0.050 | 0.0755 |
| $\rho$ | Migration proportion | 0.0333 | 0.042 |
| $\gamma$ | Migration distribution parameter | 0.01 | 0.00868 |
| init *s* | Prop. susceptible bats in initially affected counties | 0.7 | 0.7 |
| init *e* | Prop. exposed bats in initially affected counties | 0.28 | 0.28 |
| init *n* | Prop. infectious bats in initially infected counties | 0.02 | 0.02 |
| init *f* | Free-living fungus in initially affected counties | 0.1 | 0.1 |
